# Rosa26 docking sites for investigating genetic circuit silencing in stem cells

**DOI:** 10.1101/575266

**Authors:** Michael Fitzgerald, Mark Livingston, Chelsea Gibbs, Tara L. Deans

## Abstract

Approaches in mammalian synthetic biology have transformed how cells can be programmed to have reliable and predictable behaviour, however, the majority of mammalian synthetic biology has been accomplished using immortalized cell lines that are easy to grow and easy to transfect. Genetic circuits that integrate into the genome of these immortalized cell lines remain functional for many generations, often for the lifetime of the cells, yet when genetic circuits are integrated into the genome of stem cells gene silencing is observed within a few generations. To investigate the reactivation of silenced genetic circuits in stem cells, the Rosa26 locus of mouse pluripotent stem cells was modified to contain docking sites for site-specific integration of genetic circuits. We show that the silencing of genetic circuits can be reversed with the addition of sodium butyrate, a histone deacetylase inhibitor. These findings demonstrate an approach to reactivate the function of genetic circuits in pluripotent stem cells to ensure robust function over many generations. Altogether, this work introduces an approach to overcome the silencing of genetic circuits in pluripotent stem cells that may enable the use of genetic circuits in pluripotent stem cells for long-term function.

## INTRODUCTION

Pluripotent stem cells have the potential to augment tissue regeneration, in addition to creating cell-specific *in vitro* diagnostics and drug screens because they are capable of self-renewing and differentiating into any cell type (1). Induced pluripotent stem (iPS) cells can be derived from mature tissue cells from individuals by expressing key transcription factors (2). However, despite advances in stem cell culture techniques, differentiation can be inefficient, laborious, expensive, or otherwise intractable (3,4). It has been proposed that programming reliable cell behaviour using approaches in synthetic biology can be used for directing cell fate decisions to enhance their therapeutic potential (1,5,6). For example, using genetic circuits to reprogram cells to facilitate precise gene regulation can be used to enhance cell differentiation outcomes for tissue engineering and regenerative medicine applications. Moreover, to capitalize on the tight gene control of genetic circuits, their stability and function in the genome is critical.

Novel genetic tools built by synthetic biologists have transformed how cells can be reprogrammed and include genetic programs to introduce switching (7–14), oscillations (15–20), logic gates (21–23), and biosensing (24–30) behaviours into cells. Assembling simple genetic parts into more complex gene circuits can reliably and predictably control cell behaviours (31,32). To date, the majority of mammalian synthetic biology has taken place in easy-to-grow and easy-to-transfect cells, derived from immortalized cell lines. These model cell lines are useful for enhancing our understanding of synthetic gene circuits that underscore the potential of synthetic biology tools, however, these cell lines may not be good predictors of the challenges that arise in stem cells (33). Plasmid and viral gene delivery systems have been shown to lose expression over weeks of cell culture, which is thought to be a consequence of epigenetic modifications of the inserted DNA (34–36). The current understanding of transgene, a gene that has been artificially introduced into the genome, silencing suggests that silencing of genes can occur through the methylation of the expression cassette and/or the formation of heterochromatin, both of which facilitate changes in gene expression; however, the circumstances that trigger these mechanisms are still being elucidated. To overcome these challenges, several strategies to mitigate the effects of transgene silencing have been used, such as the inclusion of universal chromatin opening elements (37,38), scaffold/matrix attachment regions (39), CG free plasmid backbones (40), minicircle DNA (41), genomic insulators (42), and targeted integration in open chromatin regions (43). These approaches can significantly lessen transgene silencing, however the effects are not universal (44,45). Currently, the impact of genetic circuit silencing is not known, and strategies to alleviate silencing have yet to be explored. Genetic circuits are distinct from plasmid and viral expression cassettes because they contain multiple genetic modules that makeup the circuits, resulting in larger DNA constructs that need to be inserted into cells (46). Since genetic circuits endow cells with tight gene control, they have been a focus for directing stem cell fate to enable both timed and tuned gene expression that match the evolving requirements of the differentiating cells as they undergo cell fate decisions (47). Altogether, reliable stem cell programming with genetic circuits will require tools that integrate into the genome, have predictable expression patterns in differentiated cells, and avoid disruption of endogenous genes and pathways.

One previously shown approach to avoid epigenetic silencing of integrated genetic circuits is to engineer genomic safe harbour sites within the genome that support the integration of genetic material to those specific genomic locations, or loci. Additionally, safe harbour loci within the genome limit transgene silencing caused by unpredictable genome interactions associated with random insertions (34,48). The Rosa26 locus in mice has been observed to facilitate the ubiquitous expression of transgenes in developing and adult tissue when inserted at this site, suggesting that transgenes are active in germ cells, in addition to the differentiated progeny of those cells (49,50). This locus has also been used in Chinese Hamster Ovarian (CHO) cells for targeted integration that demonstrates stable integration (51). Therefore, the Rosa26 locus is an ideal location to target for inserting genetic circuits to study their stability and function, in stem cells.

Here, genetic circuit silencing was studied by observing circuit function in pluripotent stem cells by engineering a mouse embryonic stem (ES) cell line with three ϕC31 docking sites on one allele of the Rosa26 locus to function as a landing pad for large genetic circuits that have a matching recombinase. The 8kb genetic circuit, LTRi (Lac Tet RNAi), was chosen because it is relatively complex in architecture and the genetic circuit permits tuneable control of gene expression (9,52,53). Enhanced green fluorescence protein (EGFP) was controlled by LTRi, LTRi_EGFP, to study the ability to reverse the silencing of gene expression once stably integrated into the genome.

## MATERIAL AND METHODS

### Design of plasmids

The homology repair template for CRISPR was Gibson assembled to include 1kb homology arms, an FRT-flanked neomycin resistance gene, 3x attP sites, and a blue fluorescent protein (BFP) expression cassette (Addgene #36086). The 1kb upstream and 1kb downstream arms were amplified from purified mouse genome from AB2.2 ES cells (ATCC #SCRC-1023) and verified by sequencing (Supplemental Figure S1F). The Cas9/gRNA plasmid was obtained from the University of Utah Mutation Generation and Detection Core (gRNA homology: TGGGCGGGAGTCTTCTGGGC). Modifications were made to the LTRi genetic switch to exchange viral promoters with non-viral promoters, namely mEF1 and rEF1 promoters from the pVItro1-msc plasmid (InvivoGen #pvitro1-mcs) and the addition of an attP docking site recognition sequence. The modified LTRi genetic switch (mLTRi) was constructed by cloning the transgene module into the vector module with DraIII and XhoI cut sites, and the individual modules were put together by Gibson assembly. mLTRi_EGFP is in process of being submitted to Addgene (addgene.org).

### Cell culture

Mouse embryonic stem (ES) cells (ATCC #SCRC-1023) were maintained in high glucose knockout DMEM (Life Technologies #10829−018) supplemented with 15% ES certified FBS (LifeTechnologies #10439024), 1% nonessential amino acids (Life Technologies #11140050), 1mM L-glutamine (Life Technologies #25030-081), 0.1 mM 2-mercaptoethanol (Life Technologies #21985023), and 200 units/mL penicillin and streptomycin. The ES cells were plated on mitomycin C treated mouse embryonic fibroblast cells (Millipore Sigma #PMEF-NL-P1) that are G418 resistant. All cell lines were grown in a humidified 5% CO_2_, 37°C incubator. The feeder cells were grown in high glucose DMEM medium supplemented with 10% FBS, 1% L-glutamine solution, and 200 units/mL penicillin and streptomycin until seeded with mouse ES cells, at which time the media conditions were as stated for the ES cells.

### CRISPR modified pluripotent stem cells

Plasmid DNA containing the repair template with three tandem attP sites was co-transfected (1:1, Jetprime VWR#89129) with a plasmid containing Rosa26-Cas9/gRNA into mouse ES cells when confluency reached 70%. The cells were selected by adding 200 μg/ml neomycin (G418 Life Technologies #10131035) to the growth medium. Resistant clones were expanded and screened for the on-site genomic edit by genomic PCR. Copy number qPCR (Power Sybr, Thermofisher #4368577) was used to determine off-site integration. The neomycin resistance gene was removed by transfecting and flow sorting (Beckon Dickenson FACSAria) the candidate cell line with a plasmid harbouring EGFP-flip recombinase and assessing the reacquisition of neomycin sensitivity.

### Docking plasmids

mLTRi-EGFP cell lines were established by co-transfection with ϕC31 integrase in the 3X-attP AB2.2 mouse ES cells. Cells that had the mLTRi-EGFP plasmid contained a neomycin resistant cassette so these transfected ES cells were selected by G418, and resistant lines were clonally expanded and further screened by inducibility with 250 μM IPTG. One of the positive clones was chosen and used for the entire study reported here.

### Quantitative PCR primer design

All primers were design with NCBI’s primer blast to have a PCR product size between 70bp and 200bp and a melting temperature between 58°C and 62°C (Table S1). The primers for known copy number (GAPDH and ZFY1) were designed to not span and exon junction because the template was genomic DNA. The primers were tested to ensure that there was no alignment with non-specific sequences. Each primer was tested using a 2x dilution of genomic DNA to ensure a single melting curve peak to ensure specific binding to only the desired location.

### Quantitative PCR

Genomic DNA was isolated from the ES cells using lysis buffer (100mMTris-Cl, 5mM EDTA, 200mM NaCl, 0.2% SDS) and proteinase K as previously reported (54). The DNA was then precipitated in C_2_H_3_NaO_2_ and isopropanol and washed in EtOH. After genomic DNA elution and quantification using a NanoDrop (ThermoFisher) and a 1 ng/μl stock was created. From the 1ng/ul stock a 2x dilution was prepared for each qPCR experiment. Each well of the qPCR reaction contained: 10ul of Power SYBR Green PCR Master Mix (ThermoFisher #4367659), 2 ul of a 3 μM forward primer that anneals to, 2ul of a 3 μM reverse primer, genomic DNA, and up to 6 ul of H_2_O. The experiments were performed in triplicates on 96 well plates and used the StepOne™ Real-Time PCR System (ThermoFisher) where a standard curve was generated for each gene. The Ct value vs. the log of amount of DNA was plotted. The slope was calculated for each gene and the unknown 3x attP was compared to the known genes. Given that two copies of GAPDH exist in the genome, and only one copy of ZFY1, which is a gene on the Y chromosome (we used male ES cells) exist in the genome, CT values and slopes were compared to the attP sample.

### Reactivation of silenced gene circuit and flow cytometry

Silenced mouse ES cells were induced with 250μM IPTG in the presence or absence of epigenetic modifying drugs at the concentrations noted in text and within the figure legends. Modifying drugs were purchased from the following: sodium butyrate (VWR #89147), 5-azacytodine (Sigma Aldrich #A2385). Cells were treated with the drug for 24 hours with and without IPTG and EGFP expression was assessed by flow cytometry using a Beckman Coulter CytoFLEX S. Flow data was analyzed using FlowJo software.

To find the maximum recovered EGFP expression from silenced cells, the cells with LTRi_EGFP stably integrated into the Rosa26 locus that stopped expressing EGFP were used to determine the amount of NaB that would recover gene expression and still maintain the health of the cells by adding 250 μM IPTG and varying the amount of NaB. For studying the activation of EGFP expression in stably transfected mouse ES cells after the exposure to NaB, silenced ES cells were grown on top of a MEF layer in a 10cm^2^ tissue culture dish for 48 hours in the absence of IPTG. After exposure to NaB for 48 hours, the cells were passed to a 24 well plate containing a fresh feeder layer of MEFs in the absence of NaB. Twenty-four hours after passing the cells and removing NaB, 250 μM IPTG was added to each of three separate wells. Twenty-four hours after adding 250 μM IPTG, the cells were collected and EGFP was quantified using flow cytometry (Day 1). Forty-eight hours after passing the cells, 250 μM of IPTG was added to each of three separate wells. Twenty-four hours after adding 250 μM IPTG, the cells were collected and EGFP was quantified using flow cytometry (Day 2). Seventy-two hours after passing the cells and removing NaB, 250 μM of IPTG was added to each of three separate wells. Twenty-four hours after adding 250 μM IPTG, the cells were collected and EGFP was quantified using flow cytometry (Day 3). Seventy-two hours after passing the cells and removing NaB, 250 μM of IPTG was added to each of three separate wells. Ninety-six hours after adding 250 μM IPTG, the cells were collected and EGFP was quantified using flow cytometry (Day 4). For studying the impact of NaB on silenced cells, and the longevity of recovering circuit function, cells were grown in 250 μM NaB alone in addition to NaB and IPTG for eight days.

## RESULTS

### Generation of ϕC31 docking sites in the Rosa26 locus of pluripotent stem cells

Clustered regularly interspaced short palindromic repeats (CRISPR)/Cas9 technology enable the targeting of specific locations within the genome. Utilizing CRISPR/Cas9 with customized guide RNAs (gRNAs) to target specific locations within the genome offers a useful method for engineering landing pads for genetic circuits at desired locations within the genome. Indeed, CRISPR/Cas9 can be used for inserting new sequences of DNA, however, the efficiency is significantly decreased when inserting DNA sequences larger than 5 kb (55). Because most genetic circuits that function in mammalian cells are larger than 5 kb, inserting a landing pad that has a site-specific recombinase can increase efficiency of integration of these larger DNA constructs into the genome when recombinase recognition sites matching the recombinase are present in the plasmid containing the DNA to be inserted (51).

To allow for targeted and efficient integration of genetic circuits into the genome of pluripotent stem cells, ϕC31 docking sites were inserted into the Rosa26 locus (Figure 1A) of mouse ES cells using CRISPR/Cas9 technology. Three tandem ϕC31 integrase *attP* sites were inserted that serve as a landing pad for larger DNA additions containing an *attB* sequence (50). To create an efficient screening method, a neomycin resistance gene flanked by *FRT* sites was added upstream of the 3x *attP* sites inside the homology arms of the repair DNA plasmid, and outside of the arms, a blue fluorescent protein (BFP) cassette was included (Figure 1B). This plasmid was co-transfected with Cas9 endonuclease and a gRNA that targets the Rosa26 locus. Adding neomycin (G418) to the media of transfected cells enables for the selection of ES colonies that have the landing pad integrated into the genome at the Rosa26 locus (Figure 1C). CRISPR/Cas9 technology has shown to have off target effects, namely whole or parts of the repair template integrate into the genome at random locations, rather than just the repair DNA flanked by the homology arms in the targeted location (56–59). To screen for off-target insertions, a BFP cassette was added outside of the homology arms in the repair DNA template (Figure 1B). Therefore, any ES colony expressing BFP would indicate that the repair plasmid was incorrectly integrated into the genome and discarded. The ES cells resistant to G418 that did not express BFP were selected and verified by PCR for the on-site insertion of *attP* sites in the Rosa26 locus with primers designed to span part of the neomycin resistance gene and the genomic DNA beyond the homology arm (Figure 1C). ES clones with an on-site edit produced a 1.6 kb amplicon (Figure 1C and Supplemental Figure S1B) that contained an XbaI site and when isolated and cut with XbaI endonuclease produced the predicted 1.1 kb and 500 bp bands (Supplemental Figure S1C). DNA sequencing was also performed and further confirmed the insertion of the repair DNA in the Rosa26 locus.

**Figure 1:**
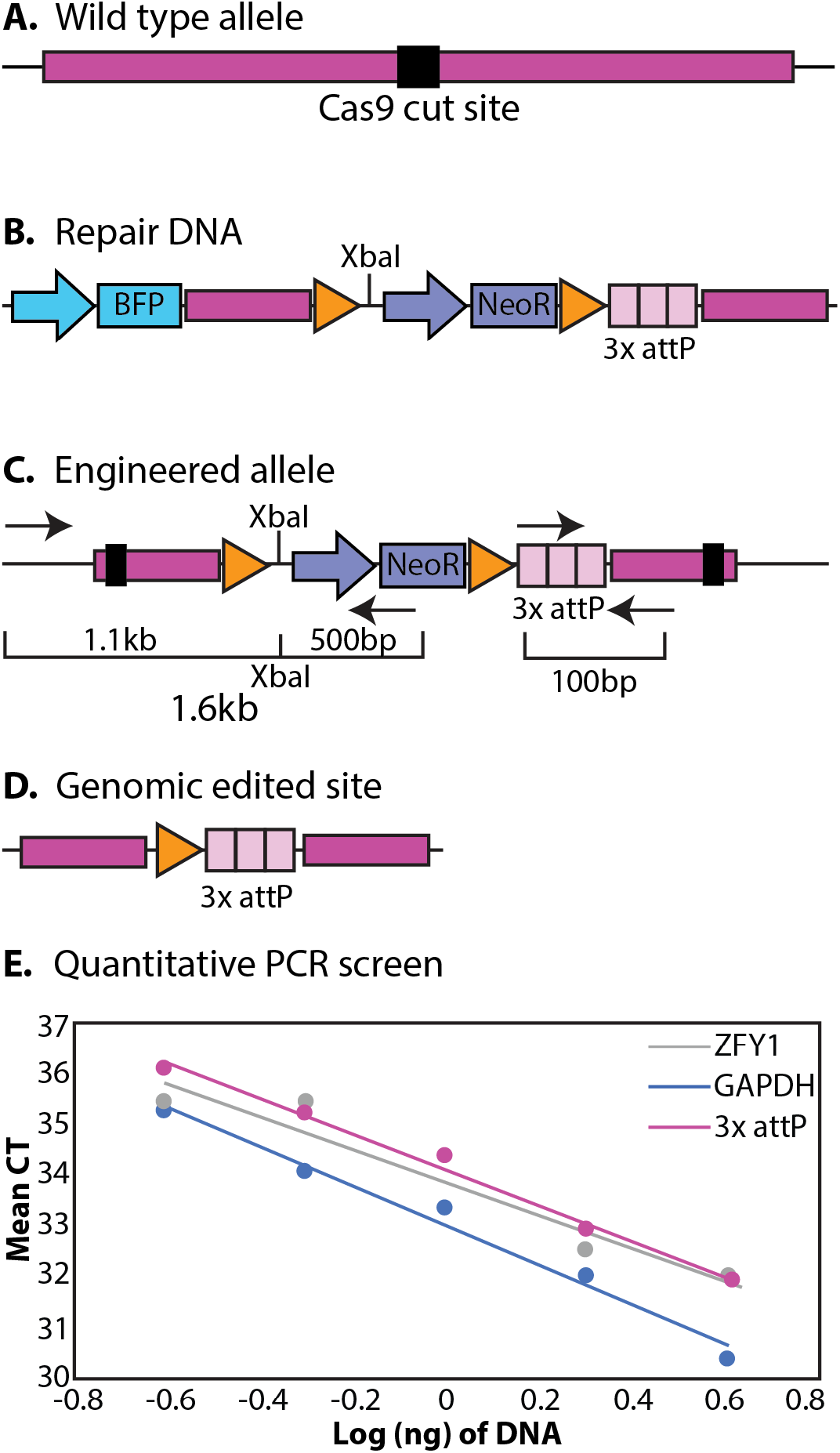
Approach for inserting docking sites into mouse pluripotent stem cells. **A.** Wild type Rosa26 allele (dark pink) with the Cas9 cut site (black square). **B**. Engineered repair DNA with 3x attP docking sites (light pink squares). Neomycin resistance (purple module) flanked by FRT sites (orange triangles) were added to enable G418 selection of the genomic insertion of the docking sites. Blue fluorescence protein (BFP, blue module) was added outside of the homology arms to screen for off-target, random integration of the repair DNA. **C**. The engineered Rosa26 allele with the repair DNA successfully inserted into the genome. The PCR screen to confirm integration into the Rosa26 locus is shown with two sets of primers (arrows). Confirmation of the NeoR module flanked by FRT sites produces a 1.6kb amplicon and confirmation of the 3x attP sites produces a 100bp amplicon. **D.** Schematic of the genomic edited site in ES clones with confirmed attP docking sites in the Rosa26 locus after transfection with flp recombinase to remove the neomycin resistance gene. **E.** Quantitative PCR on ZFY1 (grey), GAPDH (blue), and the targeted Rosa26 allele with 3x attP sites (pink).

### Screen of the mouse genome for correct genomic edits

To assess the number of docking sites integrated into the mouse genome, quantitative PCR (qPCR) was performed to verify whether additional integrations occurred elsewhere within the genome (Figure 1E). By referencing two genes of known copy number, ZFY1 on the Y chromosome (one copy), and GAPDH on chromosome 6 (two copies), it was possible to determine how many times the docking site was inserted into the genome. Primers were designed to anneal at the end of the 3x *attP* sequence and included part of the genomic Rosa26 sequence to give a ~100mer amplicon (Figure 1C). The mean Ct values of each primer set vs. the log of DNA (ng) were plotted as previously described (54). The slope of Ct values vs. log of DNA of each sample was calculated and compared to the slope of ZFY1 and GAPDH. Since the slope of the attP amplicon matched that of the ZFY1, it shows that the docking site was added once (Figure 1C). Once correct docking was confirmed, the neomycin resistance gene was removed by transfecting the confirmed heterozygous ES line with a plasmid harbouring flip recombinase, leaving the 3x *attP* sites (Figure 1D). To validate that the resistant cassette was removed, the *FLP* transfected ES cells were clonally expanded, assessed for neomycin sensitivity, and verified by PCR. The amplification of the wild type ES Rosa26 locus is expected to be 1.1kb, while the addition of the *attP* sites without neomycin is expected to increase in size to 1.4kb (Supplemental Figure S1E). Results indicate that the ES cell line has the Rosa26 site-specific addition of 3x *attP* sites at one allele and is nowhere else in the genome.

### Docking genetic circuits

To assess the function of LTRi_EGFP in mouse ES cells over time, the genetic circuit was integrated into the Rosa26 locus at the docking sites. First, to rule out the possibility of CMV and RSV being silenced (60), LTRi was modified to replace the original CMV and RSV promoters with the non-viral promoters mouse Elongation Factor-1 (mEF1), and rat Elongation Factor-1 (rEF1) (Figure 2A-B). Site-specific docking of the modified LTRi (mLTRi)_EGFP genetic circuit was accomplished by adding an *attB* sequence to the plasmid. Along with the genetic circuit, the plasmid also contained a neomycin resistance cassette for G418 selection of transfected ES cells. The mLTRi genetic circuit containing an *attB* sequence was co-transfected with ϕC31 integrase, resulting in the mLTRi genetic circuit stably integrated into the genome. ES cells resistant to G418 were clonally expanded and screened for their response to the chemical inducer, isopropyl β-D-1-thiogalactopyranoside (IPTG), a small molecule that activates gene expression of the genetic circuit (9,52,53). To confirm docking, the IPTG responsive cell lines were screened for integration into Rosa26 by conventional PCR with primers flanking the Rosa locus and the genetic circuit, which is expected to produce a 1.3kb amplicon (Supplementary Figure S2A and S2B).

**Figure 2:**
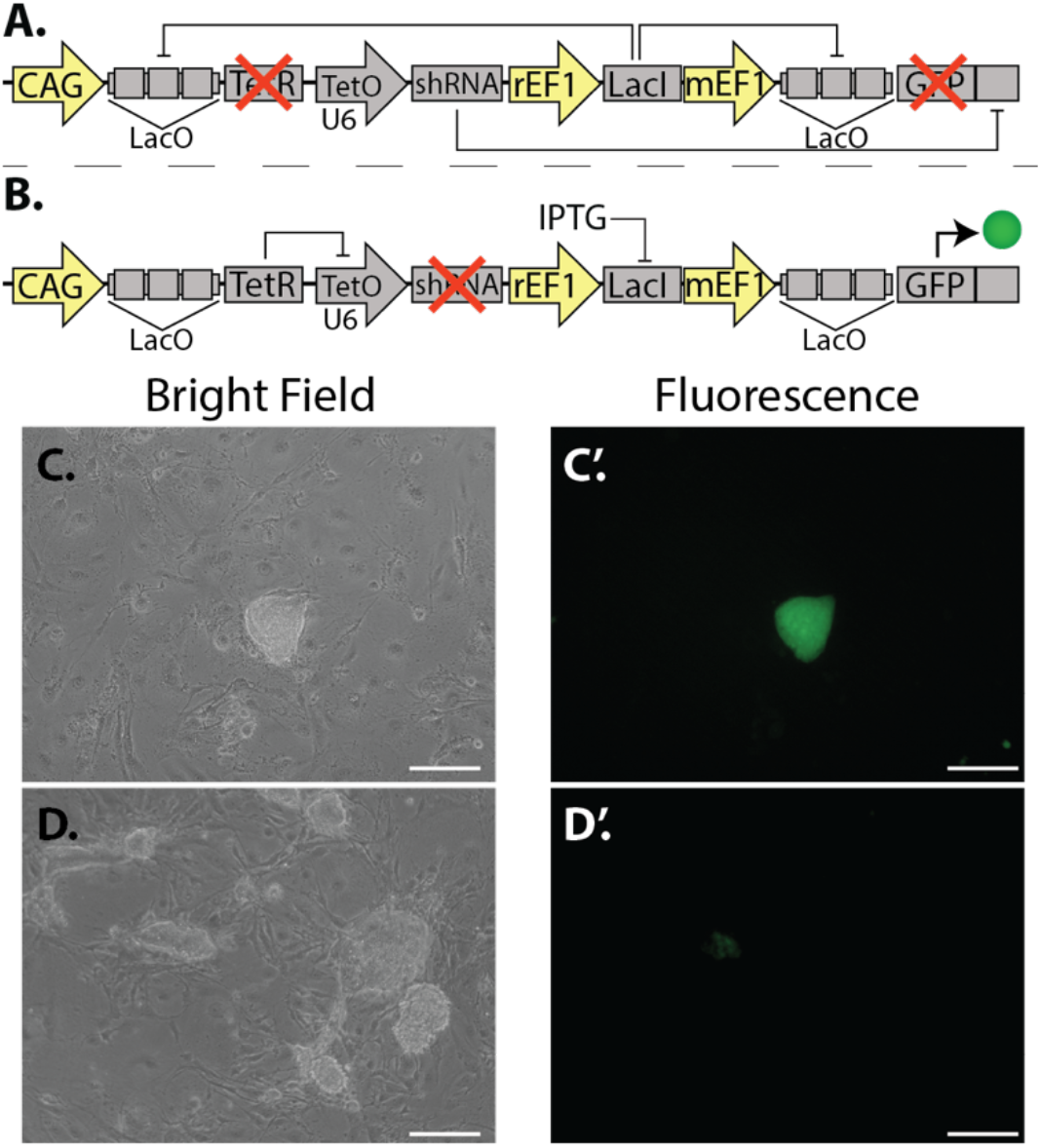
Integrating mLTRi_EGFP into the Rosa26 locus. **A.** Schematic of mLTRi in the off state: LacI repressor proteins are constitutively expressed and bind to the lac operator sites upstream of TetR and GFP. This causes transcriptional repression of TetR and GFP. With the repression of TetR, shRNA is transcribed by the U6 promoter and complementarily binds to the synthetic target sequence located in the 3’UTR of the GFP mRNA. **B.** Schematic of mLTRi in the on state: in the induced state, IPTG binds to the LacI proteins, producing a conformational change in the repressor proteins. This causes them to no longer bind to the lac operator sites, which allows for the transcription of GFP and TetR. The Tet repressor proteins bind to the tet operator site located in the U6 promoter, repressing the transcription of the shRNA. The resulting effect is a robust expression of GFP. **C.** Bright field image of stably integrated mLTRi into the Rosa26 locus of mouse ES cells 7 days after selection with G418. **C’**. The presence of 250μM IPTG in the culture media induces robust expression of EGFP 7 days after G418 selection. **D**. Bright field image of stably integrated mLTRi 30 days after selection with G418. **D’.** Fluorescence image of stably integrated mLTRi 30 days after selection with G418. Scale bars, 200μm.

### Assessing circuit function in the Rosa26 locus over time

To investigate the function of the mLTRi genetic circuit in pluripotent stem cells, mLTRi was docked into the Rosa26 locus of mouse ES cells and the cells expressing EGFP in the presence of IPTG. The EGFP expressing cells were single cell sorted and cultured for more than three weeks in the presence or absence of IPTG. EGFP was assessed using fluorescent microscopy and results show that ES colonies cultured beyond three weeks significantly lost their ability to express EGFP in the presence of IPTG compared to cells in earlier time points (Figure 2C-D’).

### Reactivation of silenced mLTRi in pluripotent stem cells

To investigate methods to reactivate mLTRi_EGFP, we looked at two common mechanisms of transgene silencing that are frequently cited as barriers when introducing transgenes were studied. The first, promoter methylation, occurs with the methylation of cytosine residues in CpG sequences by the cytosine DNA methyltransferase (DNMT1) enzyme and can be reversed with methyltransferase inhibitor 5-aza-deoxycytidine (AzaC) (61–66). The second, histone acetylation, is a process where an acetyl group is added to a lysine residue on the tail of a histone by histone deacetylase, which can be prevented by sodium butyrate (NaB), a known histone deacetylase inhibitor (HDACi) (67–69).

To test whether the silenced genetic circuit could be reactivated, ES colonies that lost their ability to respond to IPTG were grown in the presence of the inhibitors AzaC or NaB for 48 hours, and EGFP expression was assessed using flow cytometry (Figure 4A). EGFP expression was reactivated in the presence of NaB. Next, to determine the optimal concentration of NaB for recovering genetic circuit function, various concentrations of NaB were added to the media of silenced ES cells in the presence of 250 μM IPTG (Figure 3B and Supplemental Figure S3). While 500 μM NaB showed the most recovery of EGFP expression, cells grown in this conditions displayed morphological changes and the cells did not appear as healthy as the cells grown in lower concentrations of NaB. The 250 μM concentration of NaB did not appear to alter the growth rate or the morphology of the ES cells over time. Therefore, the 250 μM concentration of NaB was used in our experiments. To determine whether EGFP expression. To determine whether EGFP expression dynamics could be recovered, ES cells that recovered their EGFP expression in the presence of 250 μM NaB were sorted by flow cytometry and grown in various concentrations of IPTG (Figure 3C) over various time points. These data show that exposure to 250 μM NaB for 48 hours reverses the silencing of mLTRi_EGFP and allows for the recovery of the genetic circuit function.

**Figure 3:**
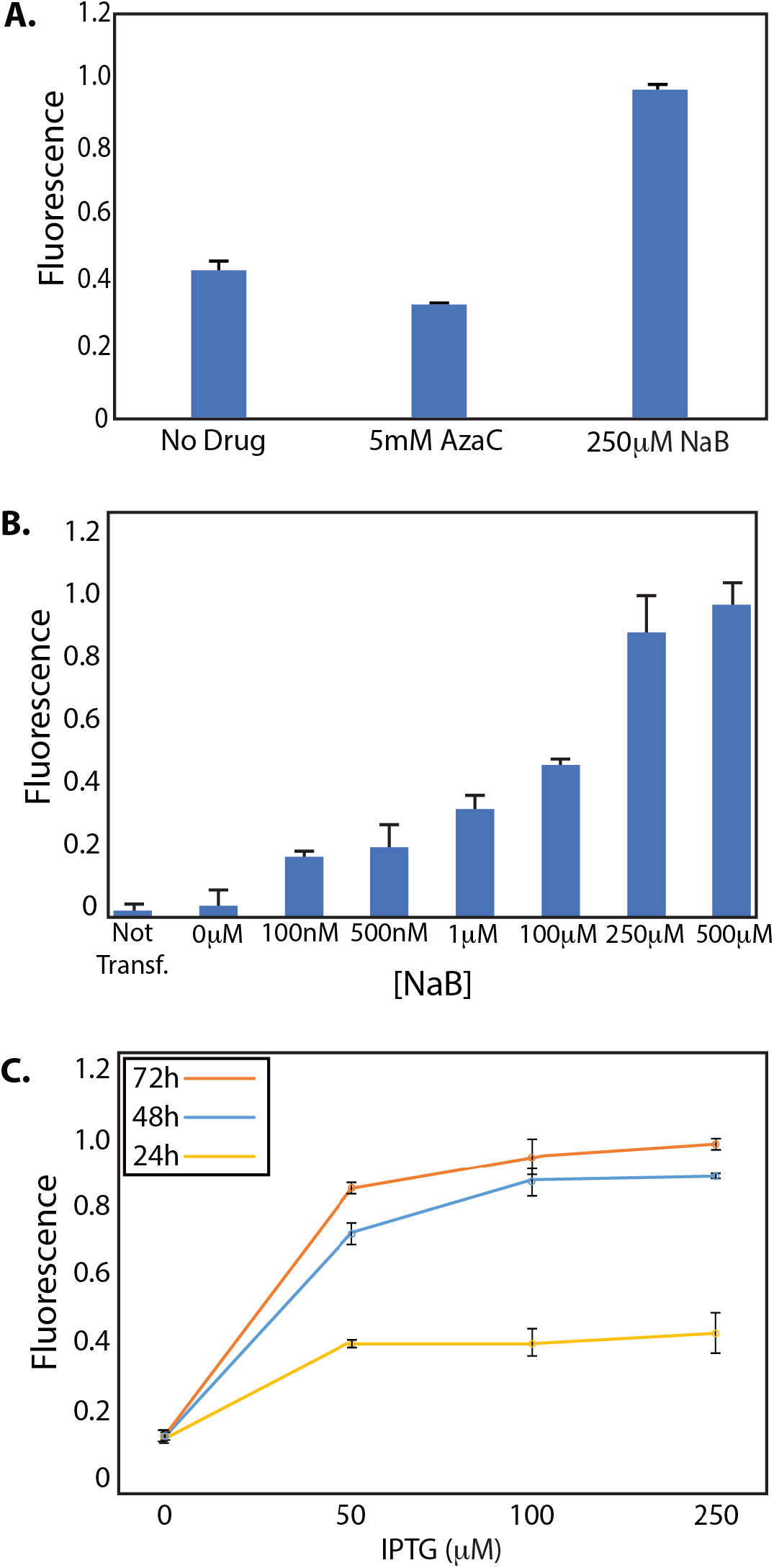
Quantification of EGFP expression after re-activation of the silenced gene circuit. **A.** Silenced ES colonies with mLTRi_EGFP integrated into the Rosa26 locus that stopped expressing EGFP in the presence of IPTG after weeks of culturing were exposed to either 5-aza-deoxycytidine (AzaC) or sodium butyrate (NaB) with 250μM IPTG in the media of each condition. EGFP expression was assessed after 48 hours of exposure to AzaC or NaB in the media using flow cytometry and the median expression levels were quantified. **B.** Median EGFP fluorescence of cells after 48 hours of growing with 250μM IPTG in the media and increasing concentrations of NaB. **C.** Median EGFP expression of recovered switching dynamics. Cultures were maintained with 250μM NaB and varying concentrations of IPTG over 24 hours (yellow), 48 hours (blue), and 72 hours (orange). In all experiments, each data point represents the median EGFP expression in at least three independent experiments. The error bars represent the standard deviation between these experiments. EGFP expression was normalized to the maximum expression in each experiment.

**Figure 4:**
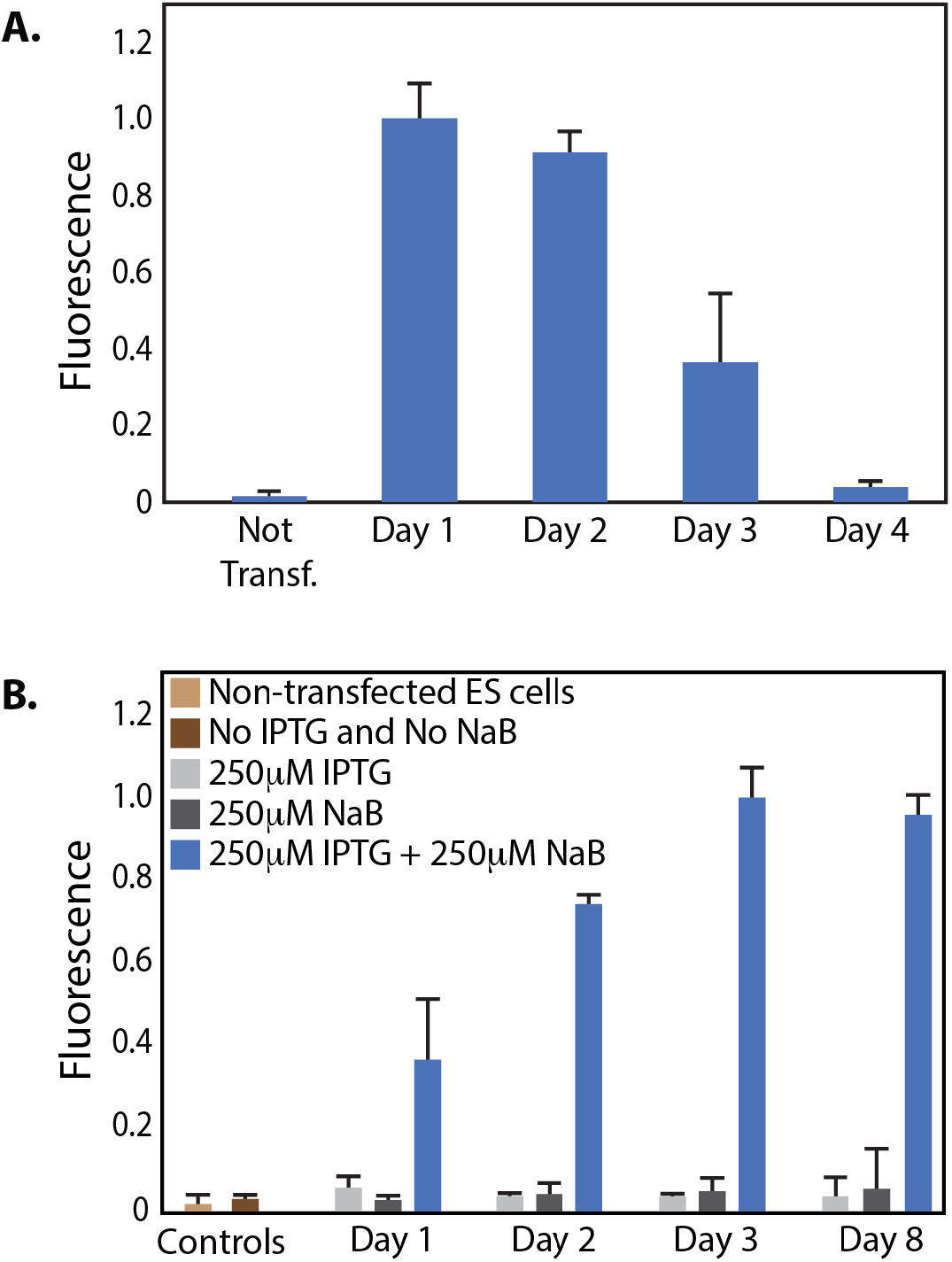
Re-activation of EGFP expression in stable ES colonies. **A.** Silenced ES cells with mLTRi_EGFP stably integrated into the Rosa26 locus and not responding to IPTG induction were grown in 250μM NaB for 48 hours then the NaB was removed. One day (Day 1), two days (Day 2), three days (Day 3) and four days (Day 4) after removal of the NaB, 250μM of IPTG was added to the media and EGFP expression was assessed using flow cytometry 24 hours after the addition of IPTG. EGFP fluorescence was normalized to the max expression, here the Day 1 data. Each data point represents the median EGFP expression in at least three independent experiments. The error bars represent the standard deviation between these experiments. **B**. ES cells with LTRi_EGFP stably integrate into the Rosa26 locus were grown in various conditions: light grey: 250μM IPTG, dark grey: 250μM NaB and blue: 250μM IPTG + 250μM NaB for up to 8 days. The light brown are non-transfected ES cells and the dark brown are ES cells with mLTRi_EGFP stably integrated into the Rosa26 locus with no IPTG or NaB added to the media. Each data point represents the median EGFP expression in at least three independent experiments. The error bars represent the standard deviation between these experiments. EGFP expression was normalized to the maximum expression.

### Investigating the activation of EGFP in stable ES cells

To explore the activation of EGFP in the silenced stable ES cell line, we first looked at inducing EGFP expression with IPTG after exposure to 250μM NaB for 48 hours (Figure 4A and Supplemental Figure S4). To determine how long after NaB exposure genetic circuits could be activated with IPTG, cells were grown in a 10cm dish in the presence of NaB for 48 hours. After 48 hours, the cells were washed and passed into 24-well plates. Twenty-four hours after passing and the removal of NaB (Figure 4A, 1 Day), IPTG was added and EGFP fluorescence was quantified 24 hours after the addition of IPTG using flow cytometry. IPTG was subsequently added 2, 3, and 4 days after the removal of NaB and the passage of cells into the 24-well plates. We observed that EGFP expression could be rescued up to 3-4 days after the removal of NaB.

To better assess whether NaB non-selectively activates gene expression, EGFP expression in stably transfected ES cells was compared to cells grown in the presence and absence of 250μM NaB and 250μM IPTG over 8 days (Figure 4B and Supplemental Figure S5). We observed that adding NaB alone to the media did not activate the expression of EGFP in the genetic circuit and that EGFP expression can be maintained for at least 8 days with the addition of NaB and IPTG.

## DISCUSSION

To date, genetic circuits in mammalian cells have primarily been reported in easy to grow and easy to transfect immortalized cell lines. Pluripotent stem cells have the potential to give rise to all cell types in the body and can propagate indefinitely under the right culturing conditions. Because pluripotent stem cells represent a single cell source that can make large contributions toward currently unmet clinical needs for regenerating damaged and diseased tissue, tightly controlling specific genes is critical for effectively driving stem cell differentiation into desired lineages. Novel genetic tools built by synthetic biologists allow for such control in a variety of mammalian cell lines. However, gene expression from transgene expression systems have been shown to have heterogenic expression patterns that are often silenced by epigenetic modifications over time (61,62). To overcome this limitation, we have engineered mouse embryonic stem cells with ϕC31 docking sites using CRISPR/Cas9 technology to allow for the targeted insertion of genetic circuits into the Rosa26 locus. This docking site functions as a landing pad for genetic circuits that have a matching recombinase to be targeted for insertion at this genome location. Docked mLTRi_EGFP in this location demonstrated robust circuit function for up to three weeks of culture, however, after three weeks, EGFP expression significantly decreased and mLTRi_EGFP was no longer responsive to IPTG induction.

To overcome silencing in pluripotent stem cells, we showed that adding the HDAC inhibitor, NaB, to the media recovers the genetic circuit function for at least 8 days. This 8-day recovery of circuit function may be sufficient for directing certain differentiation pathways that turn on early in development and/or for activating transcriptional cascades capable of regulating later cell fate signals responsible for developmental regulation (70).

Taken together, this study demonstrates that genetic circuits can be inserted into the genome of pluripotent stem cells and if circuit function diminishes over time, NaB can be added to the growth media to re-establish circuit function. These data raise the exciting possibility of using synthetic biology in pluripotent stem cells for many therapeutic applications.

## MATERIALS AND DATA AVAILABILITY

Any data or unique materials (e.g. DNA sequences) presented in the manuscript may be available from the authors upon reasonable request and may require a materials transfer agreement. The mLTRi_EGFP plasmid is currently being deposited to Addgene (addgene.org).

## SUPPLEMENTARY DATA

Supplementary data are available at SYNBIO online.

## ACKNOWLEDGEMENTS

We thank the members of the Deans Laboratory for helpful discussions on the manuscript, and the University of Utah Flow Cytometry Facility that is funded by the National Cancer Institute [5P30CA042014-24], and by the National Centre for Research of the National Institutes of Health [1S10RR026802-01].

## FUNDING

This work was supported by the University of Utah start-up funds, the National Science Foundation CAREER Program [CBET-1554017 to TLD], the Office of Naval Research Young Investigator Program [N00014-16-1-3012 to TLD], and the National Institute of Health Trailblazer Award [1R21EB025413-01 to TLD]. This work was also supported by the University of Utah Undergraduate Research Opportunities Program (UROP) to CG.

## CONFLICT OF INTEREST

None.

## AUTHOR CONTRIBUTIONS

MF constructed the mLTRi genetic circuit and made the stable cell lines. CG designed the quantitative PCR screen. ML helped with quantifying the reactivation of the genetic circuit using flow cytometry. All authors analyzed the data, wrote, and edited the manuscript.

## Supplementary Information

**Supplemental Figure S1:**
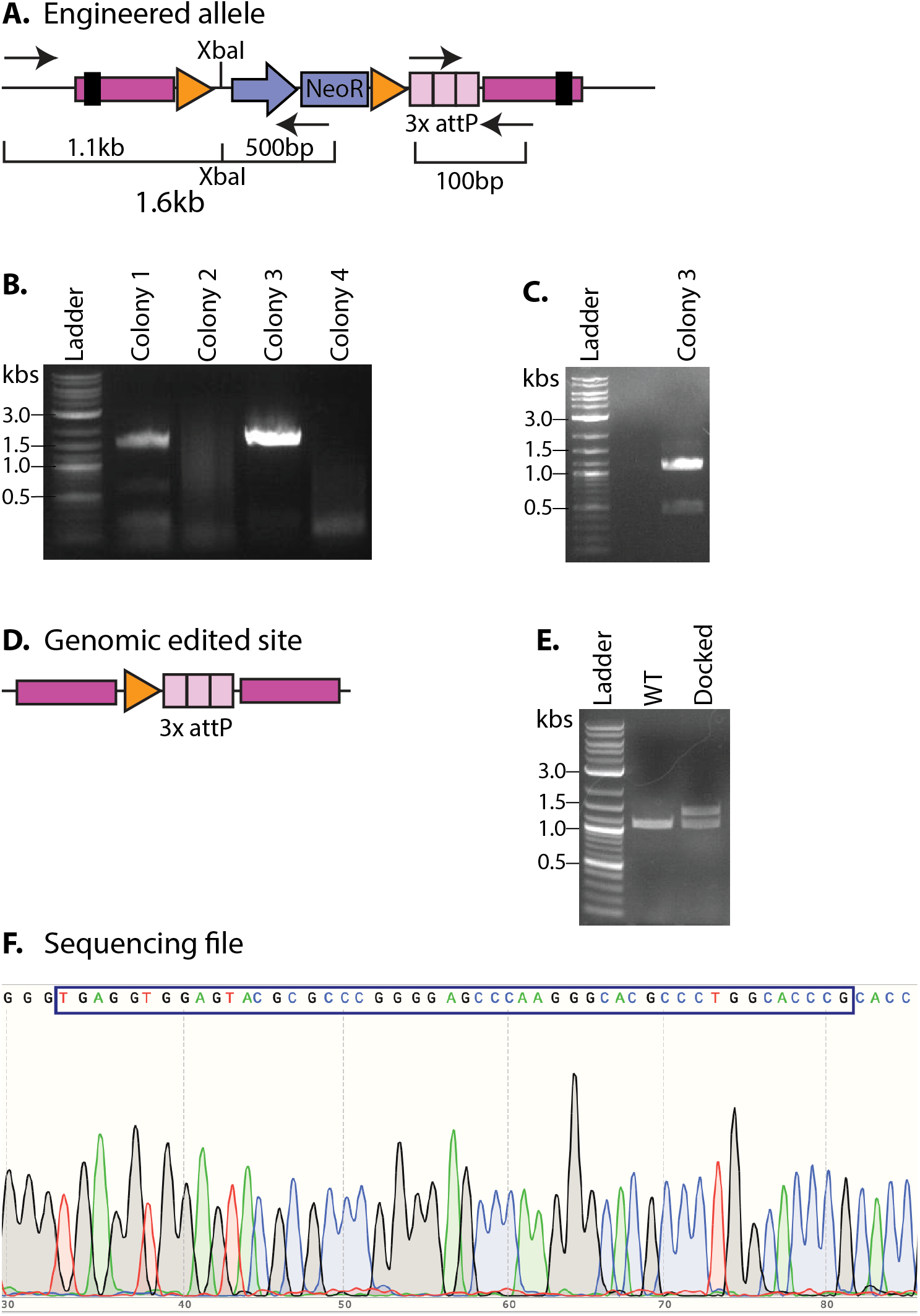
Screening for docking site integration. **A**. Schematic of PCR screen to confirm integration into the Rosa26 locus. Correct integration yields an expected amplicon size of 1.6kb. **B**. PCR on the genomic DNA of four different ES colonies. **C**. Colony 3 amplicon was gel isolated and cut with XbaI to confirm integration of the repair DNA. **D.** Schematic of the genomic edited site in ES clones with confirmed attP docking sites in the Rosa26 locus after transfection with flp recombinase to remove the neomycin resistance gene. **E.** PCR comparing the wild time (WT) ES cells without the docking site and with a confirmed ES colony harboring the 3x attP docking site. **F.** Sanger sequencing of the attB site (sequence in box).

**Supplemental Figure S2:**
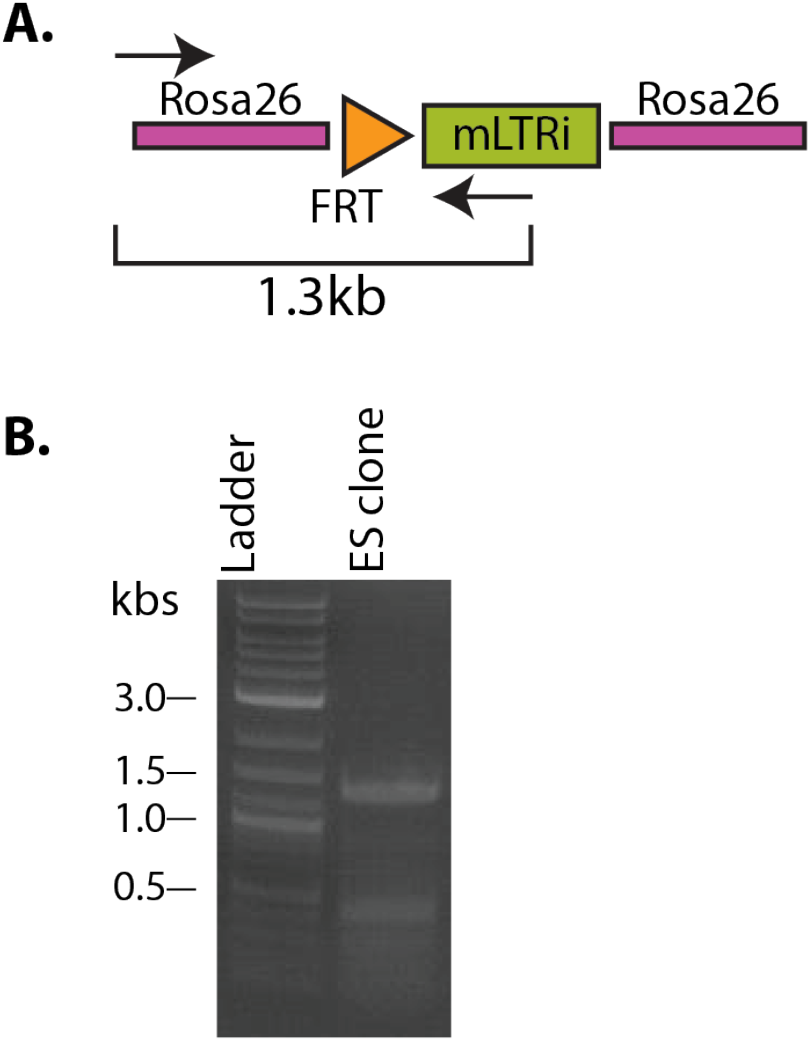
Screening docked mLTRi genetic circuit. **A**. Schematic of primer design for PCR of genomic DNA with mLTRi integrated into the Rosa26 docking site. **B.** PCR results confirming on-site integration with primers that span the genome and part of the inserted genetic circuit.

**Supplemental Figure S3:**
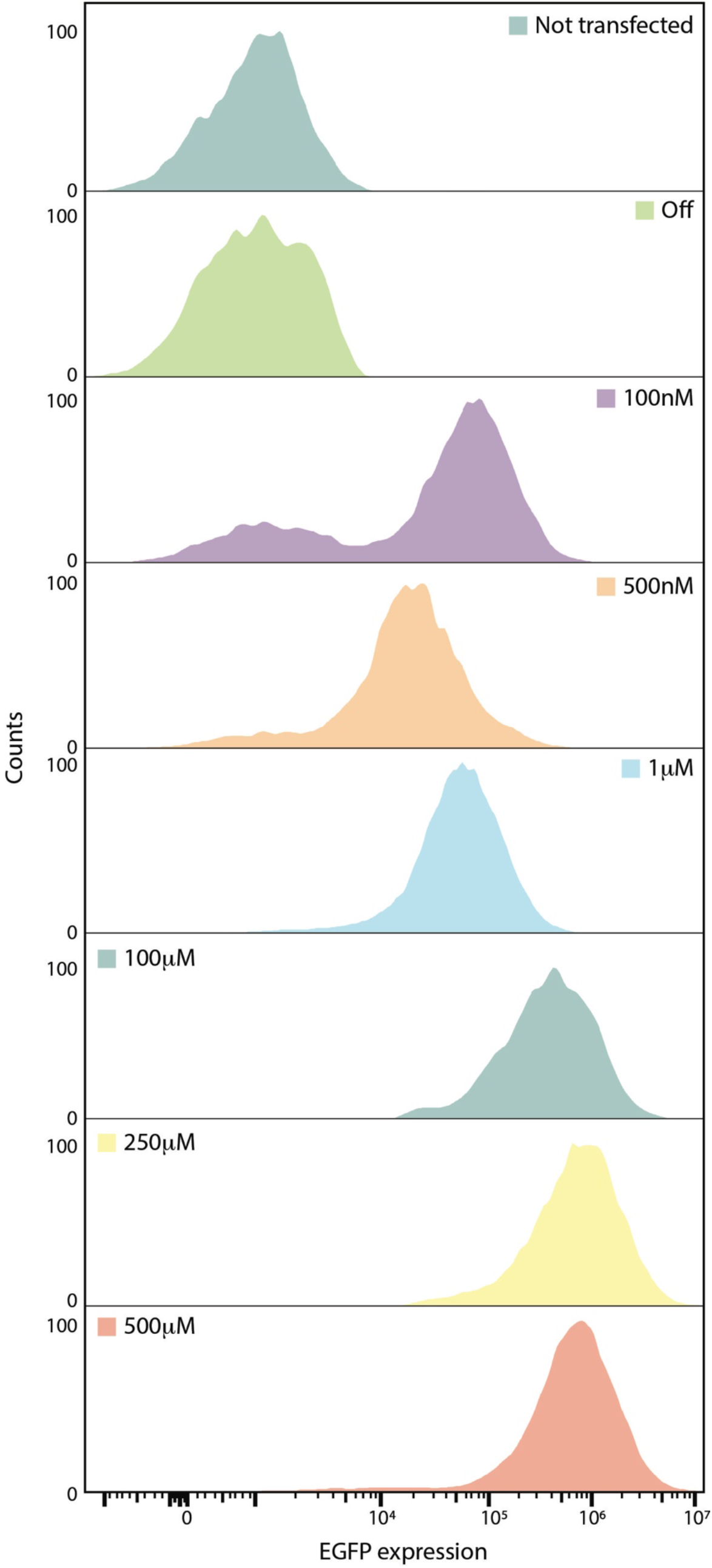
Cell population effects with varying amounts of NaB. Silenced ES cells with mLTRi_EGFP integrated into the Rosa26 locus that stopped expressing EGFP in the presence of IPTG after weeks of culturing were exposed to 250μM IPTG and varying amounts of NaB (shown in the figure) for 48 hours. After 48 hours, each population was run on flow cytometry to quantify EGFP and to assess the cell population shifts in the presence of IPTG and NaB.

**Supplemental Figure S4:**
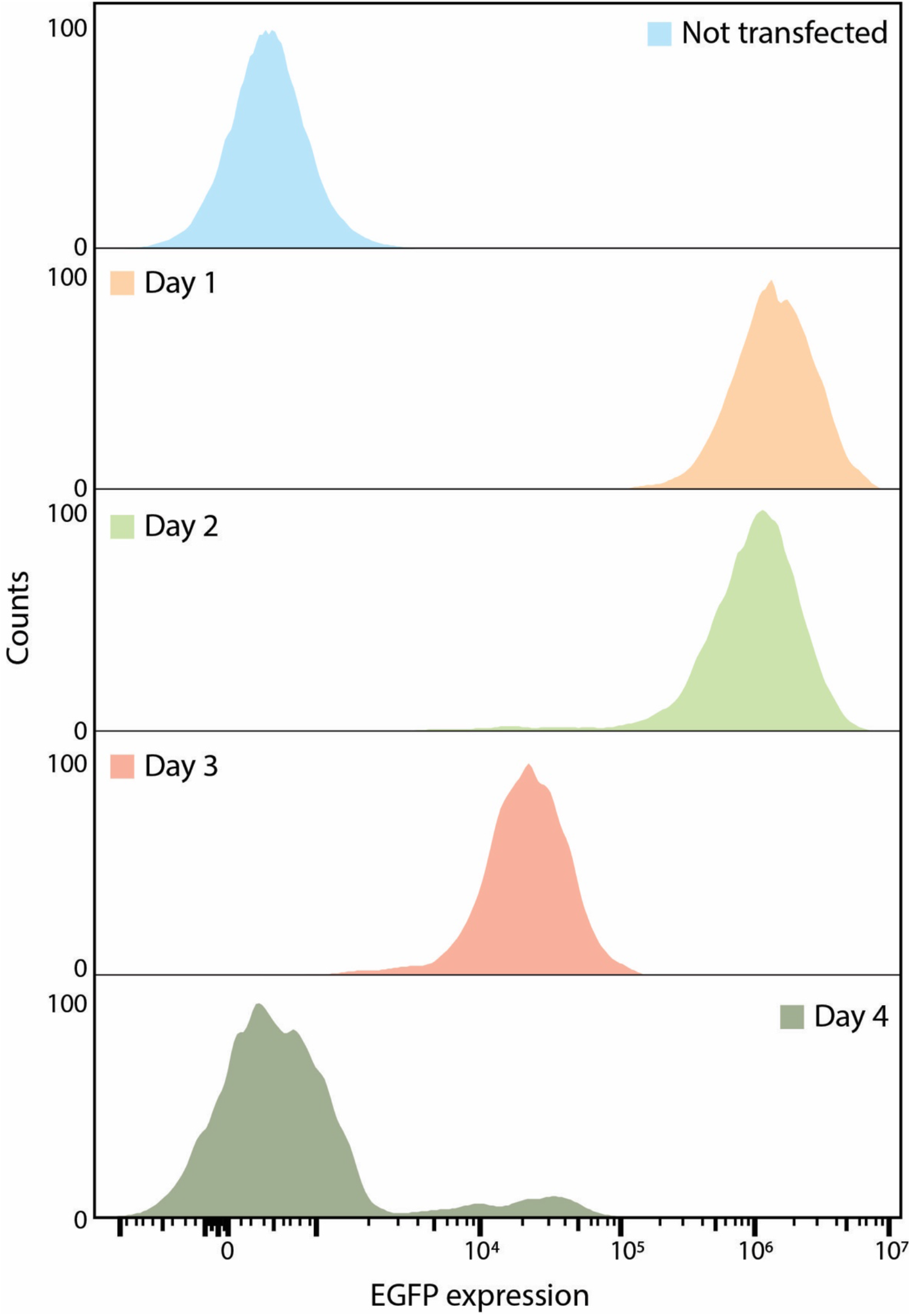
Cell population effects re-activating EGFP after removal of NaB. Silenced ES cells with mLTRi_EGFP integrated into the Rosa26 locus that stopped expressing EGFP in the presence of IPTG after weeks of culturing were grown in 250μM NaB for 48 hours then the NaB was removed. One day (Day 1), two days (Day 2), three days (Day 3) and four days (Day 4) after removal of the NaB, 250μM of IPTG was added to the media and EGFP expression was assessed using flow cytometry 24 hours after the addition of IPTG.

**Supplemental Figure S5:**
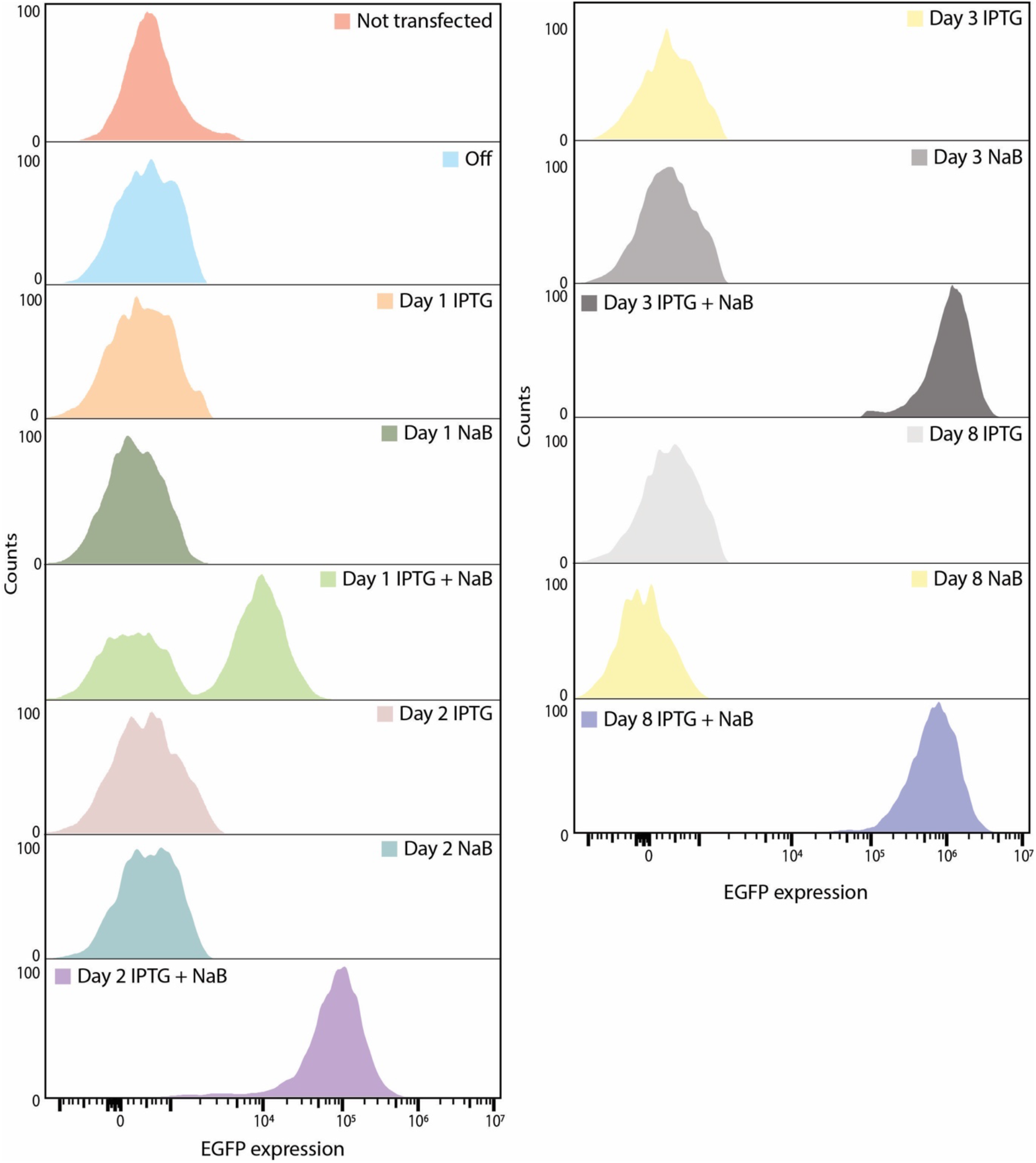
Cell population effects investigating the impact of NaB on cells. Silenced ES cells with LTRi_EGFP stably integrate into the Rosa26 locus that stopped expressing EGFP in the presence of IPTG after weeks of culturing were grown in various conditions as indicated in the figure for up to 8 days.

**Table S1:**
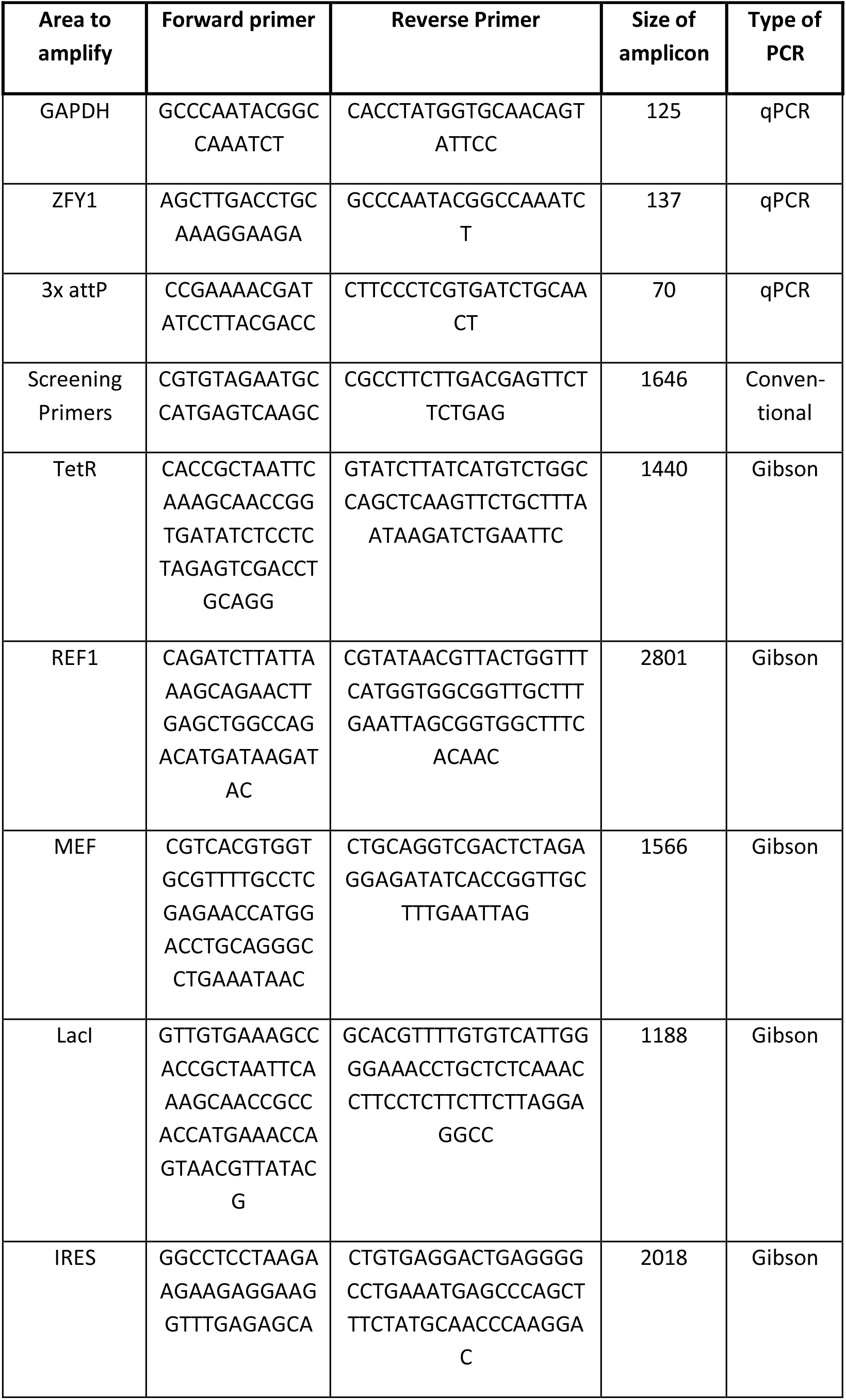
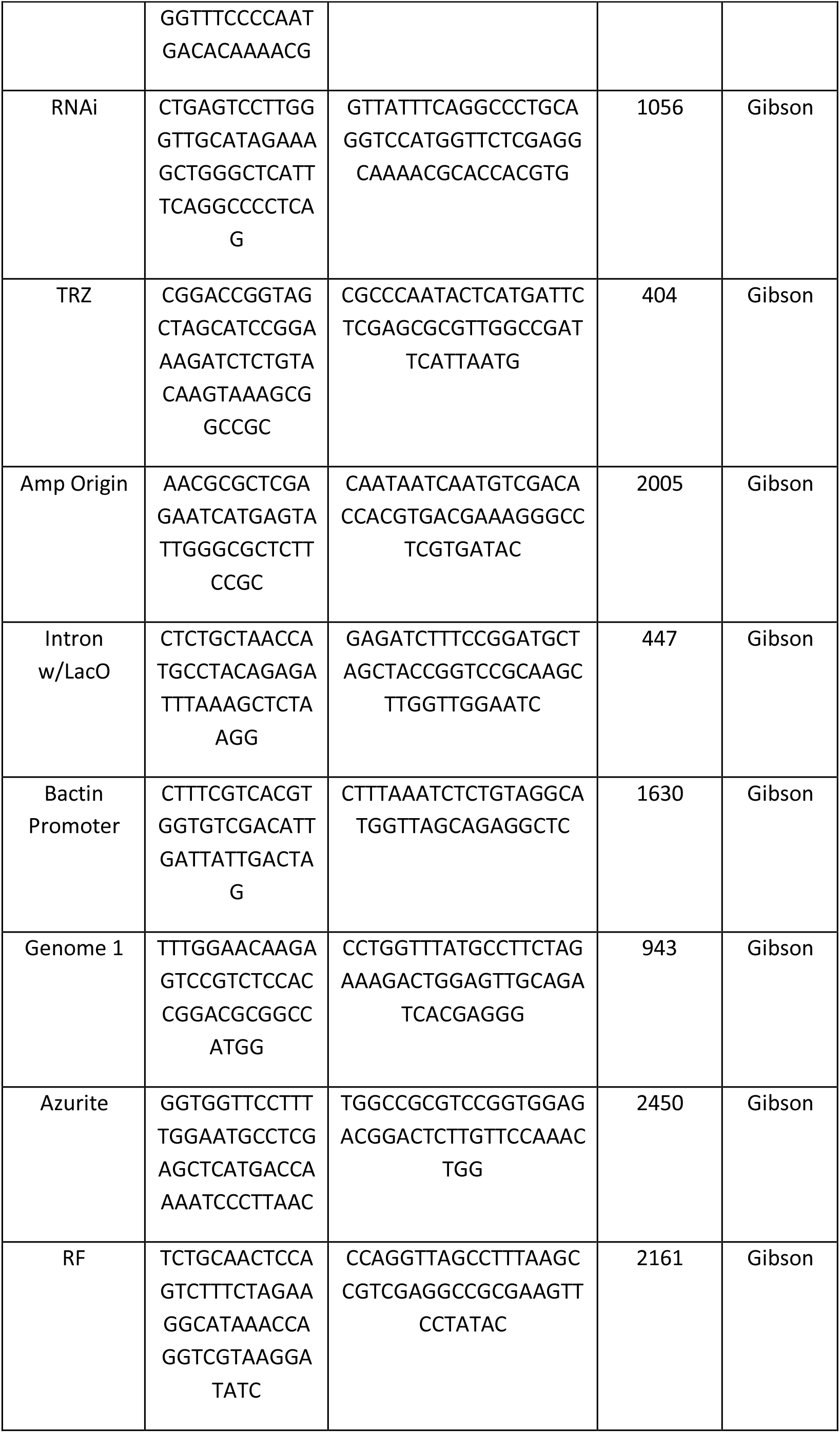
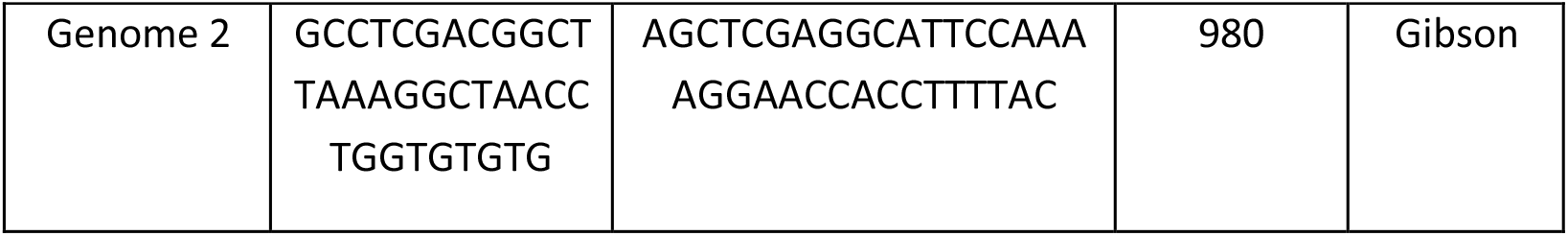
Table of primers used in the study.

### Features in mLTRi_EGFP

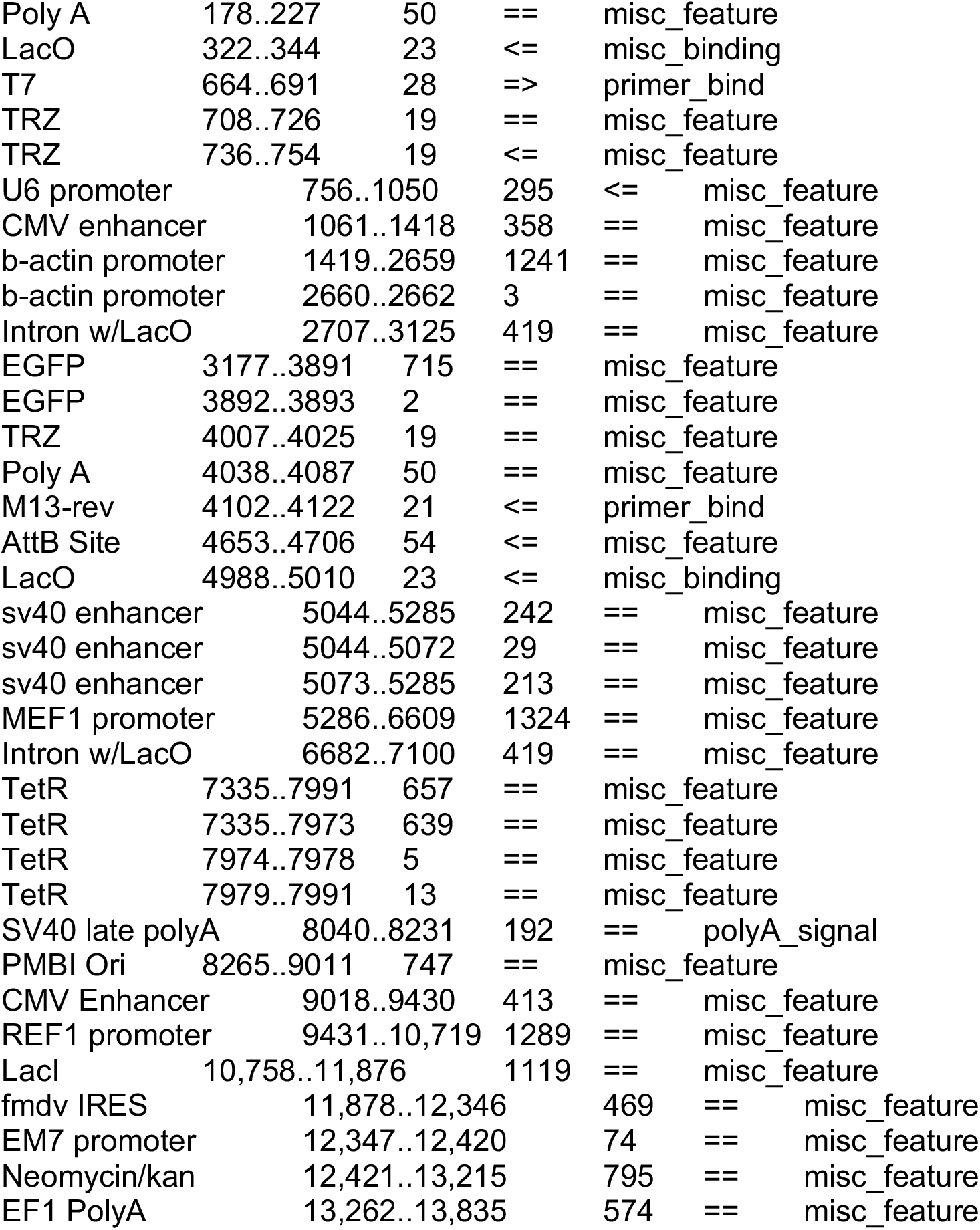

### Map of mLTRi_EGFP

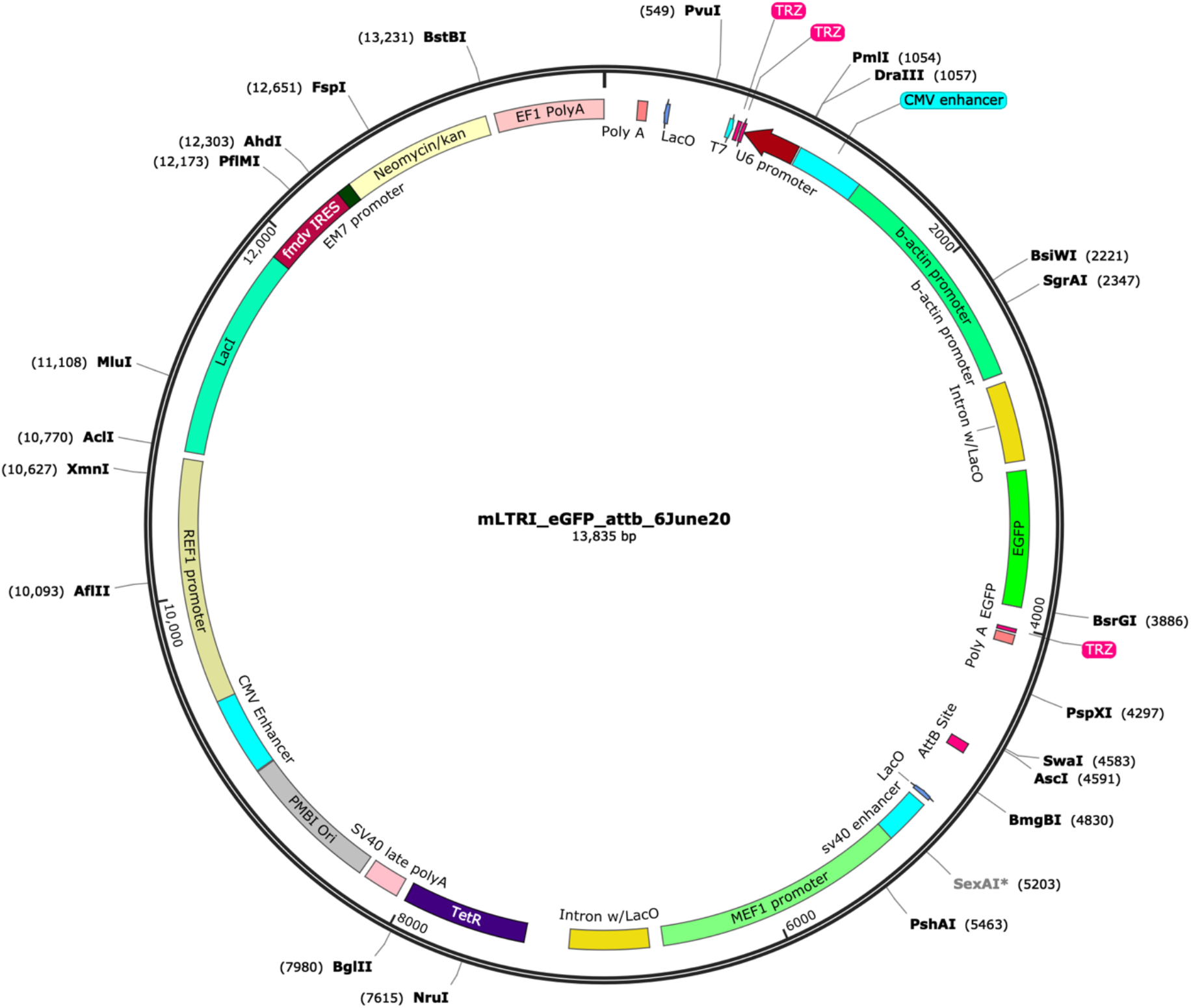

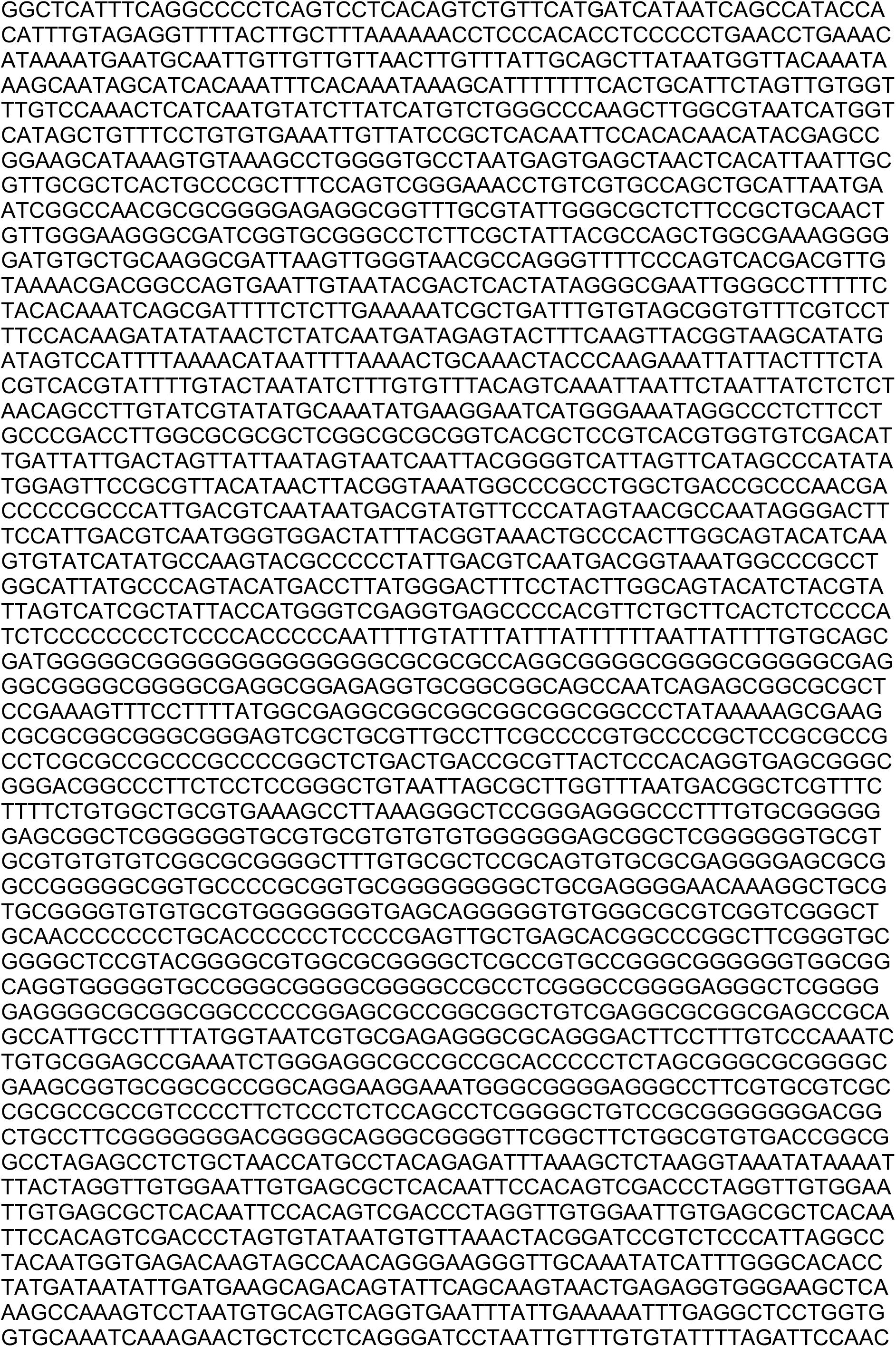
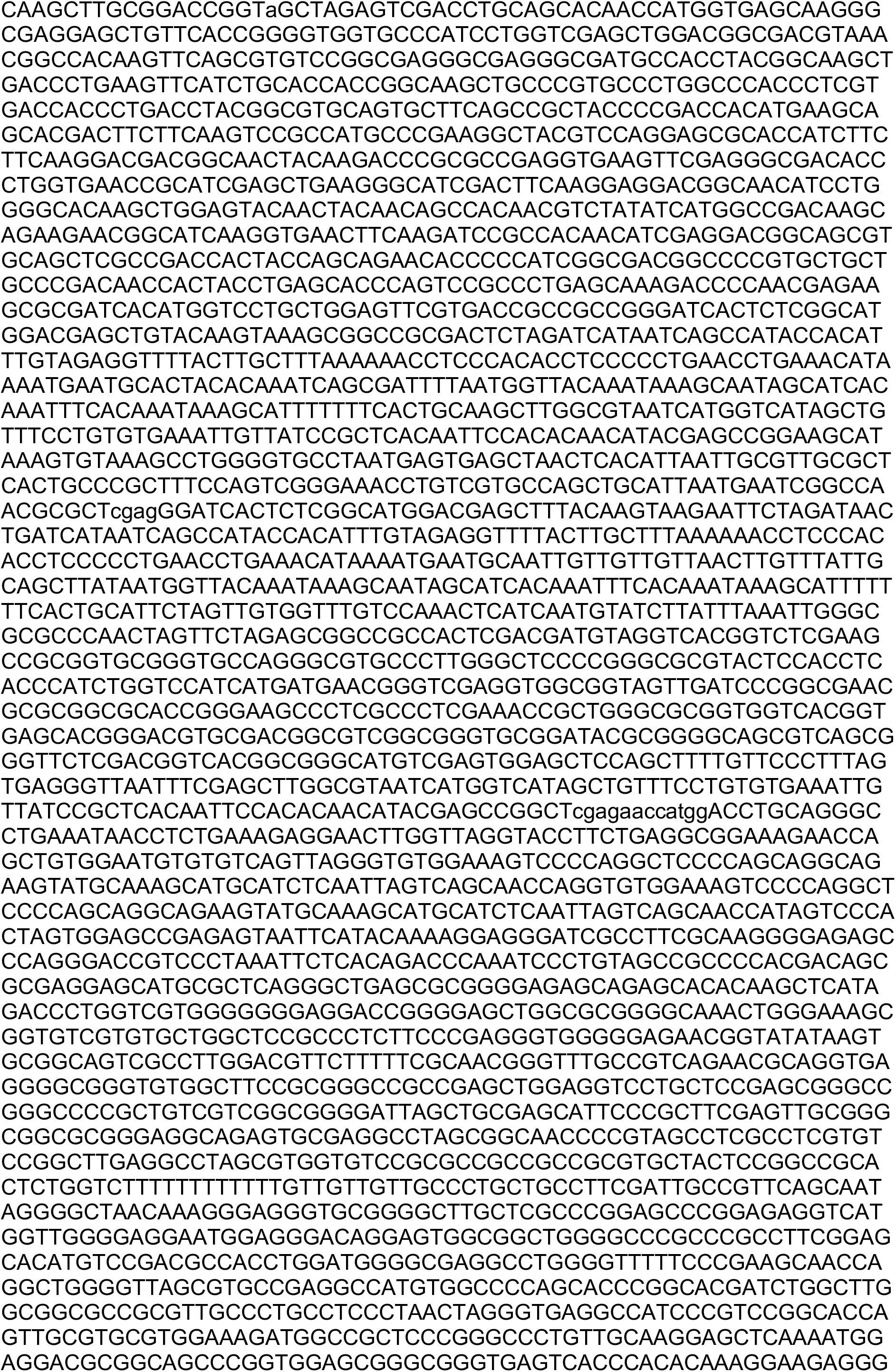
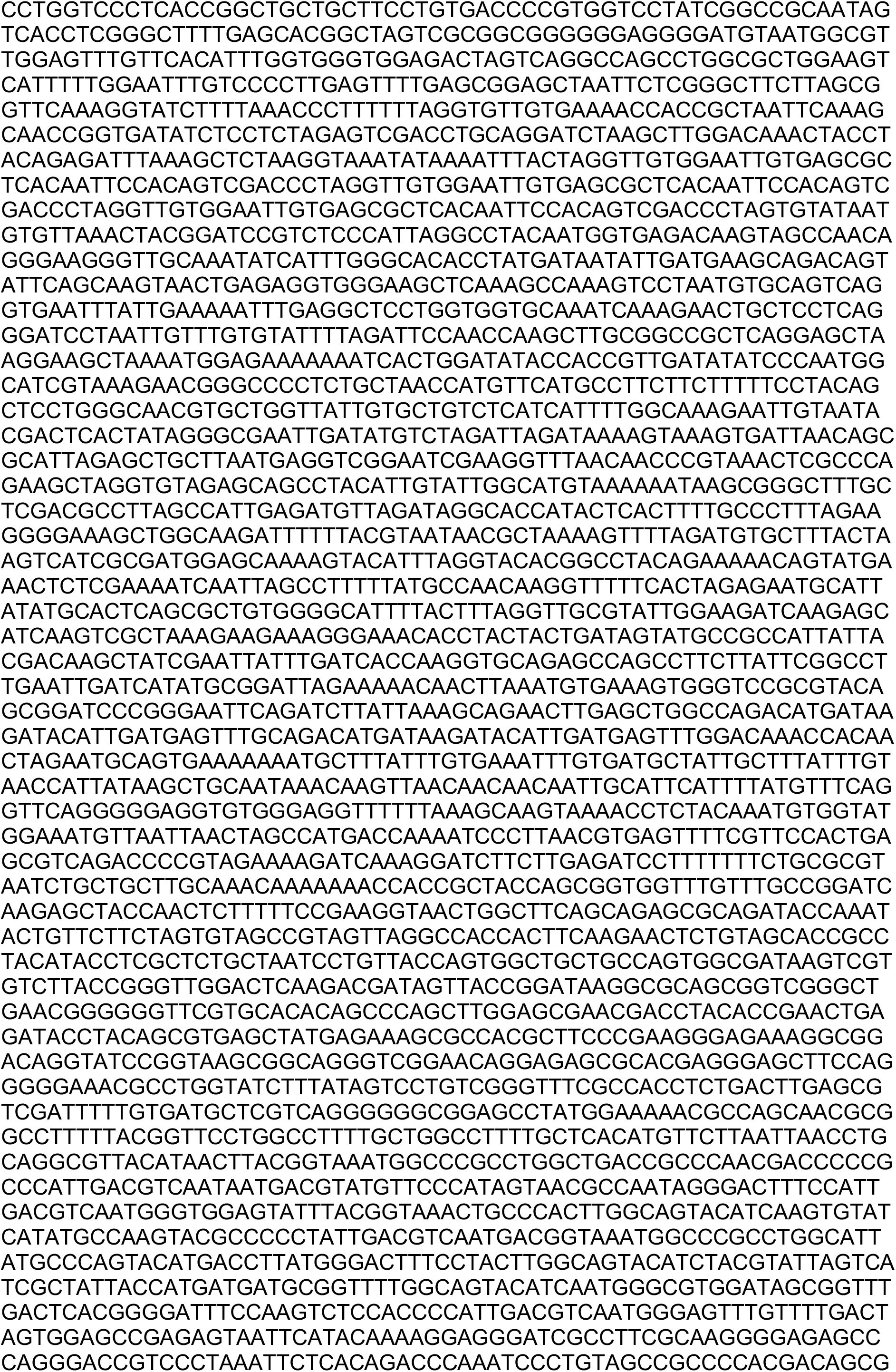
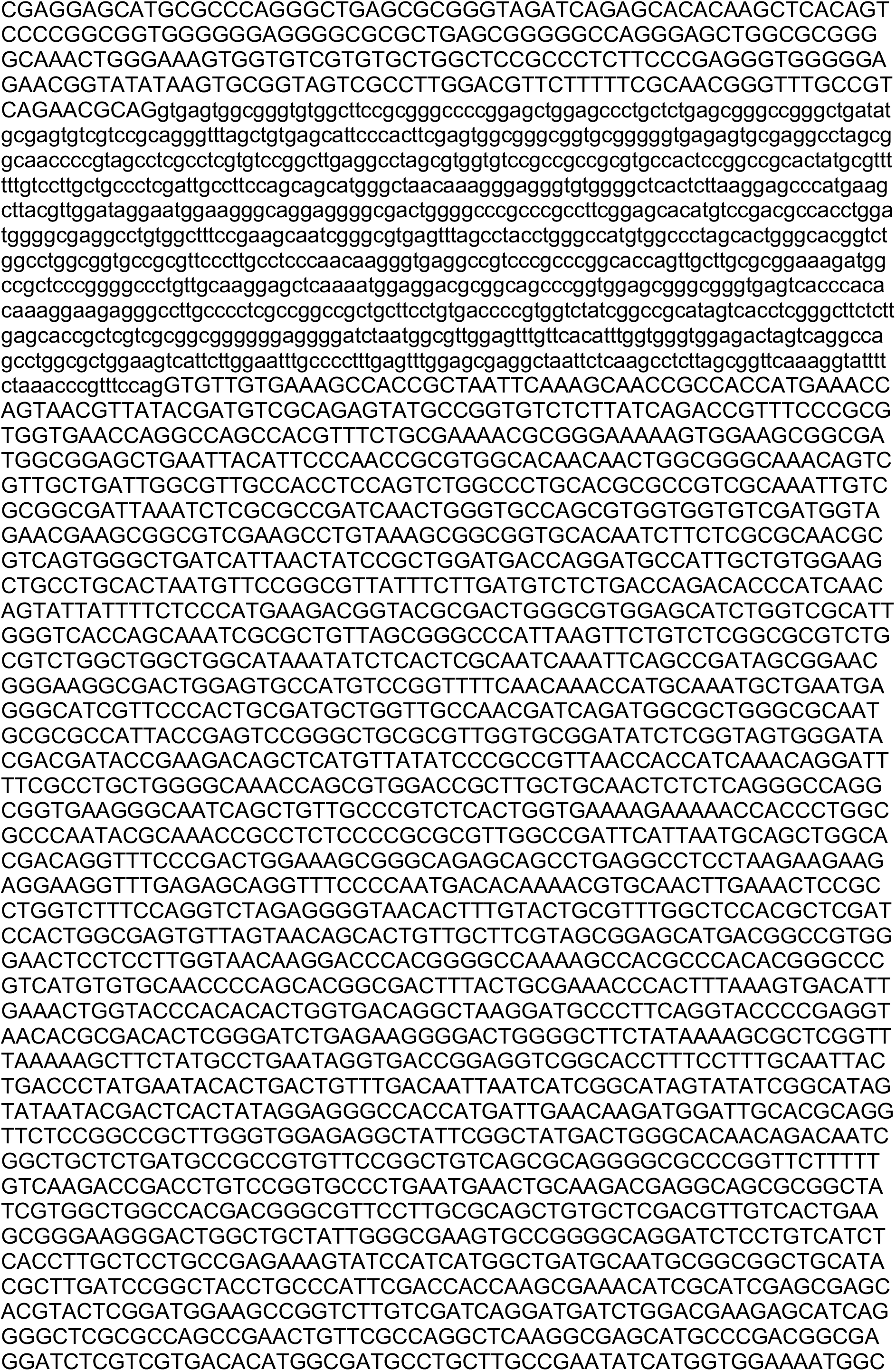
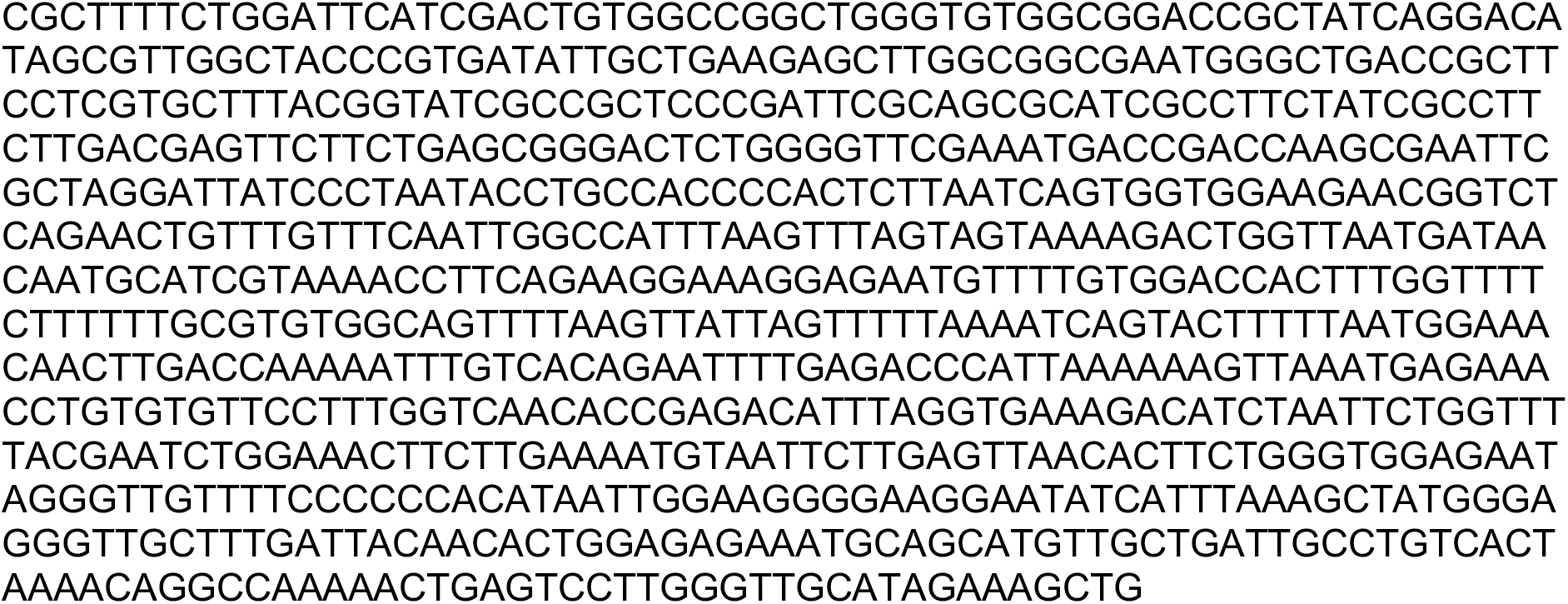

## Notes

### Competing Interest Statement

The authors have declared no competing interest.

